# Contrast-FEL – a test for differences in selective pressures at individual sites among clades and sets of branches

**DOI:** 10.1101/2020.05.13.091363

**Authors:** Sergei L. Kosakovsky Pond, Sadie R Wisotsky, Ananias Escalante, Brittany Rife Magalis, Steven Weaver

## Abstract

A number of evolutionary hypotheses can be tested by comparing selective pressures among sets of branches in a phylogenetic tree. When the question of interest is to identify specific sites within genes that may be evolving differently, a common approach is to perform separate analyses on subsets of sequences, and compare parameter estimates in a *post hoc* fashion. This approach is statistically suboptimal, and not always applicable. Here, we develop a simple extension of a popular fixed effects likelihood method in the context of codon-based evolutionary phylogenetic maximum likelihood testing, Contrast-FEL. It is suitable for identifying individual alignment sites where any among the *K ≥* 2 sets of branches in a phylogenetic tree have detectably different *dN/dS* ratios, indicative of different selective regimes. Using extensive simulations, we show that Contrast-FEL delivers good power, exceeding 90% for sufficiently large differences, while maintaining tight control over false positive rates. We conclude by applying Contrast-FEL to data from five previously published studies spanning a diverse range of organisms and focusing on different evolutionary questions.

## Introduction

When the same gene is subjected to different selective environments, population processes, or exogenous adaptive forces in different sets of species or other taxonomic units (e.g., viral or bacterial isolates or cancer lineages), especially if it leads to functional adaptation or differentiation, we expect to find distinct molecular signatures of selection among these sets. At the nucleotide or protein level, this difference can manifest as variation in evolutionary rates across groups of species, for example in the *rbcL* gene of monocots where species with shorter generation times showed higher evolutionary rates (Gaut et al., 1992), or both across sites and lineages, leading to *heterotachy* – a process that is widespread in protein evolution and has been studied extensively (e.g., Lopez, Casane and Philippe (2002); Whelan, Blackburne and Spencer (2011)).

At the codon level, a commonly adopted modeling framework is to allow the strength of selection, represented by the ratio of non-synonymous (*dN*) and synonymous (*dS*) substitution rates, *ω*:= *dN/dS*, to vary across branches or both branches and sites. The primary focus of methodological development has been to estimate *ω* and compare it to the neutral expectation *ω*:= 1 (e.g., see Delport, Scheffler and Seoighe (2008) or Arenas (2015) for a review). Here we focus instead on the methods for comparing *ω* across sets of branches; These methods are relatively few and far-between (cf Table 1). We further assume that the branches are partitioned into groups using additional sources of information, and not inferred as a part of the evolutionary analysis. Yang (1998) developed a likelihood ratio test to compare *gene-average* selective pressures among different sets of branches in the tree. By design, Yang’s method relies on pooling data across sites and branches, and lacks resolution to identify individual sites subject to selective differentials. A more recent model allowing clade-level effects in a site-mixture framework can infer fractions of sites experiencing a clade-level shift, but not individual sites that differ between branches (Baker et al., 2016). Another approach allows *ω* to vary across sites and branches as a random effect, with the group-level effect to distinguish the sets of branches (Wertheim et al., 2015). This method can answer more refined questions (e.g., Is selection on one set of branches relaxed or intensified relative to the other set?), but it still lacks the site-level resolution. Several approaches have been specifically designed to detect evidence of directional selection towards a preferred subset of residues at specific sites and/or branches in a phylogenetic tree. Parto and Lartillot (2017) developed random effects mutation-selection models that allow selective profiles of amino acids to vary across sites and branches and can be fitted in a Bayesian phylogenetic framework; they can identify specific sites and specific residues subject to directional selection. Tamuri, dos Reis and Goldstein (2012) implemented a conceptually similar model in a fixed effects model and fitted it using maximum likelihood approaches to identify which residues are preferred at specific sites. Murrell et al. (2012) augmented standard codon models to reward/penalize substitutions towards specific residues along predefined sets of branches at specific sites (using fixed effects) and applied it to the evolution of drug resistance in HIV. Rife Magalis et al. (in press) applied a version of Contrast-FEL to determine which HIV-1 envelope sites might be evolving differentially between three anatomical compartments in a single host.

**Table 1.** Methods for comparing selection among different sets of branches. ML: maximum likelihood, LRT: likelihood ratio test, MCMC: Markov Chain Monte Carlo

| Method | Application | Statistical Framework |
| --- | --- | --- |
| Yang (1998) | Gene-wide differences in average $\omega$ | ML, LRT |
| Baker et al. (2016) | Gene-wide differences in distributions of $\omega$ | Random Effects ML, LRT |
| Wertheim et al. (2015) | Gene-wide differences in distributions of $\omega$ | Random Effects ML, LRT |
| Tamuri, dos Reis and Goldstein (2012) | Site level amino-acid preferences | Fixed Effects ML |
| Murrell et al. (2012) | Site level directional selection | Fixed Effects ML, LRT |
| Parto and Lartillot (2017) | Site and branch-level amino-acid preferences<br>$\omega$ variation at site / branch level | Bayesian MCMC Mixture |
| Contrast-FEL | Site-level differences in $\omega$ | Fixed Effects ML, LRT |

Here we fully develop and validate a fixed effects site-level model (Contrast-FEL) and a likelihood ratio test to formally test the hypothesis of differences in *ω* ratios between two or more groups of branches using a likelihood ratio test. We evaluate Contrast-FEL using comprehensive simulations, and, having established reasonable statistical behavior, apply it to five disparate empirical datasets previously analyzed for differential selection among branch sets.

## Methods

### Statistical models and parameter inference

Our model adapts the Fixed Effects Likelihood (FEL) method (Kosakovsky Pond and Frost, 2005), which has previously been used to find sites that evolve under different selective pressures in different alignments of the same gene (Kosakovsky Pond et al., 2006). Consider an alignment of *N* coding nucleotide sequences with *S* codons, and a given tree topology 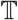 with *B ≥ N* branches. Branches in the tree are partitioned into disjoint non-empty sets, *B_i_, i* = 1,2*,…,K ≥* 2, and this partition is fixed *a priori*. If *K >* 2, one of the branch groups can be designated as background and not explicitly tested. Sequence evolution is modeled using the Muse-Gaut class of codon models, MG-REV-CF34, with the general time reversible component for the nucleotide substitution process and the *CF* 3*×*4 estimator for the equilibrium frequencies (e.g., see Murrell et al. (2013)). The key parameters of the model are: *α* – the site-specific synonymous substitution rate, and *β_i_* – the site specific non-synonymous substitution rate for branch group *i*, and all others are nuisance parameters. We fit the model to a coding sequence alignment using the following procedure.

Step 1 Obtain initial nuisance parameter estimates: infer *B* branch length parameters *t_b_*, and 5 nucleotide substitution rates *θ_AC_,θ_AT_,θ_CG_,θ_CT_,θ_GT_* using the general time reversible nucleotide model.
Step 2 Infer initial codon tree scaling and group-level *dN/dS*: fit the MG-REV-CF34 model to the entire alignment with each branch group having its own *dN/dS* parameter, using *C ×*nucleotide(*t_b_*) for branch lengths (*C* is the estimated tree scaling constant), and *θ* estimates from the previous step.
Step 3 Refine nuisance parameters and group-level *dN/dS*: remove constraints from branch lengths and *θ* then perform a maximum likelihood fit under the MG-REV-CF34 model.
Step 4 For each site (independently), infer parameters of interest. Fix nuisance parameters at estimates from the previous step. Estimate *α* as tree-wide branch-length scaler, and estimate *β_i_* for each group class. The site-model fitted here becomes the universal alternative hypothesis (most general model) for all site-level tests. The empirical validity of such estimation procedures has been discussed in Scheffler, Murrell and Kosakovsky Pond (2014). As a computational shortcut, invariable sites are skipped, because maximum likelihood parameter estimates are 0 at such sites.

### Hypothesis testing

Depending on the value of *K* and whether or not the background set is present, different testing procedures will be carried out at each site. All tests are likelihood ratio tests, using the assumed asymptotic distribution of 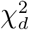 to test significance. The degrees of freedom parameter, *d*, varies from test to test.

1. *K* = 2. The single null hypothesis, *β*_1_ = *β*_2_, is tested against the universal alternative with *d* = 1.
2. *K >* 2, no background. An omnibus test using the null *β*_1_ = *β*_2_ = *… β_K_*, vs the universal alternative, with *d* = *K −*1. In addition, all pairs of groups are tested for equality of rates using the null *β_i_* = *β_j_* for 1 *≤ i< j ≤ K* with *d* = 1, resulting in *K*(*K −*1)*/*2 tests.
3. *K >* 2, with background. Without loss of generality, assume that the background is designated as group *K*. An omnibus test compares the null *β*_1_ = *β*_2_ = *… β_K_ −*1, against the universal alternative, with *d* = *K −*2. In addition, if *K >* 3, all pairs of groups are tested for equality of rates using the null *β_i_* = *β_j_* for 1 *≤ i< j ≤ K −*1 with *d* = 1, resulting in (*K −*1)(*K −*2)*/*2 tests.

When multiple hypotheses are tested at each site, the corresponding p-values are corrected to maintain nominal family-wise error rates using the Holm-Bonferroni (Holm, 1979) procedure.

### Permutation tests

As an option, it is possible to perform branch set permutation tests at each site where some of the LRT tests from the previous section are significant at a given level (e.g., *p ≤* 0.05) to assess whether or not the differences detected across groups are due to “outlier” effects. To do so, we randomly shuffle branch assignments to sets (maintaining the number and size of the sets), and perform the complete LRT procedure described above for each permuted branch set, up to 20 times. If we find the LRT as high or higher as observed on the original partition for **any** of the tests, at iteration *j*, we report permutation p-value of 1*/j* (and stop the process), otherwise we report a permutation p-value of 0.05.

### Final reports

For each site that is not invariable, Contrast-FEL outputs Holm-Bonferroni corrected p-values for each of the conducted tests, (if selected) the overall permutation p-value, and q-values (false discovery rates) computed from the omnibus test p-values using the Benjamini and Hochberg (1995) procedure.

### Simulated data

To evaluate Contrast-FEL statistical and predictive performance, we simulated 3706 data sets comprising a total of 1594400 codon sites. Simulations were designed to cover a range of relevant parameter values.

- **Tree topologies**. We simulated data along a single, empirical tree topology with reported branch lengths from Yokoyama et al. (2008) containing *N* = 31 taxa, chosen to represent a somewhat typical use case for these types of studies. We also simulated data from multiple random, balanced (maximally symmetric) and ladder-like (maximally asymmetric) topologies with *N* from 8 to 256 sequences (drawn uniformly using latin hypercube sampling [LHC], see below).
- **Branch lengths**. For each parametrically generated topology, we drew the mean branch length uniformly from 0.001 to 0.25 using LHC, and then generated branch lengths from the exponential distribution with this mean.
- **Alignment length**. Integer alignment length was drawn (uniformly) from the 100 to 800 range using LHC.
- **Branch sets**. We used several simulations where the branches from the Yokoyama et al. (2008) tree were partitioned into two groups by hand. For parametrically simulated topologies, we selected the fraction of branches to belong to the “test” set from 0.01 to 0.8 using LHC, and the rest of the branches were in the “reference” set. We also simulated 458 datasets where branches were partitioned into 4 groups.
- **Fraction of sites with different selective regimes**. The proportion of sites in an alignment evolving under the null hypothesis (*β*_1_ = *β*_2_ = *…β_N_*) was drawn for each dataset from the beta distribution with parameters *p* = 7*, q* = 1. There were also 300 strict null simulations, i.e., simulations where all sites were evolving under the null hypothesis.
- **Site to site rate variation**. Synonymous rates were either constant across sites or varied according to the gamma distribution with parameters *α* = *β ∼* max(0.1*,θ*), where *θ* was a draw from the normal distribution *N* (1,0.5). Non-synonymous rates were drawn from the gamma distribution with parameters *α* = max(0.1*,N* (0.5,0.25))*,β* = 1.

Latin hypercube sampling was performed over the set of four parameters: number of leaves, mean branch length, alignment length, and fraction of branches in the “test” set. To generate *S* LHC samples of parameter values, each parameter range is divided into *S* equiprobable intervals, and a joint sample of multiple parameters is chosen to ensure that every single interval for any parameter is sampled exactly once. LHC allows one to sample a broad range of parameter values with a relatively small number of samples.

### Implementation

The maximum likelihood estimation procedure consists of the following steps and is implemented in HyPhy v2.5.2 or later (Kosakovsky Pond et al., 2019). Steps 1-3 benefit from multi-core processors via likelihood function parallelization, and step 4 can be distributed to multiple worker nodes via MPI, improving performance. Documentation on how to use and interpret the analyses and prepare data and trees for submission to Contrast-FEL is available and at `hyphy.org.` A version of Contrast-FEL will be maintained on the Datamonkey web service (`datamonkey.org`, Weaver et al. (2018)).

### Sequence aligments

Empirical alignments and phylogenetic trees used for analysis here can be downloaded from data.hyphy.org/web/contrast-FEL/ in NEXUS format. Simulated datasets and simulation parameters can be downloaded from the same URL. Additional information is included in the `README.md` file.

### Computational performance

Following the initial model fits, site-level tests are embarrassingly parallel, and can take full advantage of distributed computing resources (MPI clusters), and scale linearly in the number of unique site patterns.

## Results

### Establishing method performance

#### False positives

The rejection rate on null data with two branch sets, i.e., on all sites where the evolutionary rates among branch sets were all equal, in aggregate, was slightly lower than nominal rates (Fig. **??**.A). We restricted the calculations only to variable sites because Contrast-FEL returns null results on invariant sites by definition (all MLEs for rate parameters are 0 at such sites) and because including invariant sites would only lower observed rejection rates. Contrast-FEL may become anti-conservative for very high divergence rates (Fig. **??**.A,E, see below). The rate of false positives on null sites was largely independent of the values of synonymous and non-synonymous rates, the levels of mean sequence divergence (within reason) in branch sets, and dataset size (Fig. **??**.B-D). The test is more conservative for low rates and smaller sets of branches, as expected. Permutation p-values and FDR q-values delivered, by design, more conservative detection rates than standard LRT p-values, but mirrored the trends of the latter (Fig. **??**.C-D). When a site is very divergent (or saturated), i.e. the product of the maximal rate estimate (*α* or *β*) and the alignment-wide tree length in expected substitutions per site exceeds 100, the test becomes anti-conservative; the q-q plot for the sites with log_10_ divergence rate between 2.5 and 3.5 is shown in Fig. **??**A. Our implementation reports the total branch length for each tested site, and saturated sites can be screened out using this metric. Such sites are rare in simulated data, and should be even more rare in empirical alignments. Only 1 in 100 simulated datasets showed false positive rates of 8% or greater (10% or greater for 1 in 1000), implying that one rarely encounters a simulated dataset where the false positive rate is notably above the nominal level.

#### Precision and Recall

The ability of Contrast-FEL to identify sites that experience differential selective pressures is influenced by the effective sample size, which depends in turn on the number of branches in the group and the extent of sequence divergence, and the effect size, i.e., the magnitude of differences in non-synonymous substitution rates, *β*. For simulations with two sets of branches, restricted to detectable sites (i.e., sites that were not invariable), power to detect differences aggregated over simulation scenarios is summarized in Table 2. Over all detectable sites, the power using using the false discovery rate of 20% is 0.319. Restricted to the sites where the differences in *β* rates between groups was at least 1 (“Large effect”), the power rises to 0.603, and further restricting to only those sites where both the test and the background branch sets had at least 3 expected substitutions per site (“Large sample size”), increases the power to 0.860. Similar trends occur for testing using LRT p-values, or permutation-based p-values. Perfectly ladder-like trees, on average yield somewhat higher power than either perfectly balanced or random/biological trees.

**Table 2.** Power of Contrast-FEL for detecting differences in selection. *N* - total number of differentially selected sites in the set. Large effect is defined as having the absolute difference in simulated *β* rates of at least 1. Large sample size is defined as having at least 3 substitutions occurring along both test and reference branch sets.

| Simulation | $N$ | $p \leq 0.05$ | $q \leq 0.2$ | Permutation $p \leq 0.05$ |
| --- | --- | --- | --- | --- |
| Overall | 139753 | 0.418 | 0.319 | 0.361 |
| Large effect | 30923 | 0.727 | 0.603 | 0.665 |
| Large effect / sample size | 14265 | 0.902 | 0.860 | 0.867 |
| Perfectly balanced trees | 39671 | 0.411 | 0.316 | 0.354 |
| Perfectly ladder-like trees | 27439 | 0.479 | 0.378 | 0.417 |
| Random/biological trees | 72643 | 0.399 | 0.298 | 0.343 |
| 4-class simulations |  |  |  |  |
| Overall (omnibus) | 18141 | 0.355 | 0.276 | 0.379 |
| Overall (any test) | 18141 | 0.415 | N/A | N/A |
| Large effect (omnibus) | 8684 | 0.516 | 0.411 | 0.542 |
| Large effect (any test) | 8684 | 0.587 | N/A | N/A |

**Table 3.** Sites evolving differentially between the Treated and the Naive sets in the HIV-1 RT dataset, at *p* ≤ 0.05. *α* – maximum likelihood estimate (MLE) of the site-specific synonymous rate; *β* – non-synonymous rate; *\** – permutation p-value is 0.05; substitutions are counted along branches in the corresponding set using joint maximum likelihood inference of ancestral states under the site-level alternative model, codons in bold are known to be involved in drug resistance (Rhee et al., 2003), ^†^ – codon identified as directionally evolving in Table 2 of (Murrell et al., 2012). FEL p-values are computed by separately testing for non-neutral evolution on the corresponding set of branches, with the + or - sign indicating the nature of selection (positive or negative). FEL pattern encodes the inferred pattern of evolution for treated/naive branches: P - positively selected (at *p* ≤ 0.05), C – conserved, N – neutral.; for example PC means “positively selected” on the treated set, and “conserved” on the naive set.

| Codon | $\alpha$ | $\beta$ (substitutions) | | p-value | q-value | Standard FEL p-value | | FEL pattern |
| --- | --- | --- | --- | --- | --- | --- | --- | --- |
|  |  | Treated | Naive |  |  | Treated | Naive |  |
| 44 | 1.31 | 0.00 (9) | 1.13 (8) | 0.0286 | 0.799 | 0.003(-) | 0.885(-) | CN |
| **65** | 1.16 | 2.12 (11) | 0.00 (3) | 0.0156* | 0.655 | 0.226(+) | 0.075(-) | NN |
| **67** | 1.24 | 1.39 (20) | 0.00 (3) | 0.0207* | 0.694 | 0.792(+) | 0.024(-) | NC |
| **70** | 1.31 | 1.56 (17) | 0.00 (5) | 0.0374* | 0.963 | 0.737(+) | 0.051(-) | NN |
| **75** | 0.86 | 1.80 (15) | 0.00 (4) | 0.0161* | 0.600 | 0.130(+) | 0.087(-) | NN |
| **100** † | 1.56 | 3.26 (29) | 0.00 (13) | 0.0150 | 0.836 | 0.094(+) | 0.075(-) | NN |
| **103** † | 1.47 | 36.51 (104) | 0.00 (7) | 0.0000* | 0.000 | 0.000(+) | 0.073(-) | PN |
| **151** † | 0.93 | 2.67 (10) | 0.00 (8) | 0.0150* | 0.719 | 0.023(+) | 0.124(-) | PN |
| **181** † | 3.32 | 4.41 (21) | 0.00 (7) | 0.0010* | 0.083 | 0.442(+) | 0.004(-) | NC |
| **184** † | 0.00 | 8.29 (58) | 0.34 (1) | 0.0000* | 0.000 | 0.023(+) | 0.110(-) | PN |
| **188** † | 0.18 | 2.99 (14) | 0.00 (0) | 0.0061 | 0.411 | 0.000(+) | 0.491(-) | PN |
| **190** † | 1.52 | 3.41 (33) | 0.00 (10) | 0.0004* | 0.041 | 0.031(+) | 0.011(-) | PC |
| **215** † | 0.44 | 1.50 (12) | 0.00 (13) | 0.0255* | 0.775 | 0.021(+) | 0.199(-) | PN |
| 228 † | 1.53 | 1.30 (21) | 0.00 (9) | 0.0436 | 0.974 | 0.753(-) | 0.019(-) | NC |
| 302 | 0.62 | 0.00 (1) | 8.05 (3) | 0.0420* | 1.000 | 0.458(-) | 0.054(+) | NN |

The power of the Contrast-FEL adheres to the expected patterns. It increases with the sample size and the effect size. For example, greater levels of divergence at a site (up to a point) corresponded to notable gains in the power of the test (Figure **??**A), as did greater numbers of substitutions occurring in the test set of branches, with power rising from *∼* 25% (*p ≤* 0.05) for 2 substitutions to 50% for 8 substitutions (Figure **??**B). Best power is achieved when the difference between substitutions rates on the two sets of branches is large (Figure **??**C), exceeding 80% for sufficiently disparate rates, and dropping to *<* 10% for rates that are very similar. Power numbers are high when the size of either of the sets is not too small (Figure **??**D).

Next, we focus on the data simulated with the relatively small (31 sequences) biological tree of vertebrate rhodopsins from Yokoyama et al. (2008) and three different test branch sets: small clade, large clade, and branches grouped by phenotype (absorption wavelength), shown in Figure **??**. For sufficiently stringent FDR (q-values) cutoffs, high (90%) precision (positive predictive value) can be achieved for all three cases, although the cutoffs need to be more stringent for the small clade scenario. High precision is achieved at the cost of fairly low recall (20*−*25%), and the small clade scenario has the worst performance among the three scenarios considered.

#### Four branch classes

Contrast-FEL remained conservative on null data when we applied it to alignments simulated with 4 branch classes (Figure **??**A), for all types of tests: FWER (family-wise error rates) corrected pairwise differences, omninbus test (any rates are different), and when considering simulations where only some (but not all) of the groups had equal rates. As was the case with simpler 2-class simulations, Type I error for severely saturated / divergent sites was somewhat elevated. Power to detect differences among any pair of branch groups, either via the pairwise or the omnibus test was strongly influenced by the effect size, ranging from near 0 for rates that were close in magnitude, to over 80% for sites where the largest substitution rate was sufficiently high (*>* 1), and sufficiently different (e.g. 5*×*) larger than the smallest rate (Figure **??**B). Power of the method is strongly influenced by the effect size, i.e., the magnitude of differences between *β* rates (Figure **??**C), and the information content/divergence of the site, measured as a function of expected substitutions per site (Figure **??**D). Introducing multiple branch classes increases the number of tests performed at each site, and because of the site-level multiple test correction, dilutes the power compared to the 2-class case (Table 2). Calling a site differentially evolving if any of the tests return a significant corrected p-value realizes a 5-6% power boost compared to relying only on the omnibus test.

If a differentially evolving site was identified as such by the omnibus test at *p ≤* 0.05 (FWER corrected), in 99.6% of cases it was also identified by one or more of the individual pairwise tests, implying that in most cases (at least for our simulation), it is possible to pinpoint specific pairs of branch sets that are responsible. For the remaining 0.4% of sites, the omnibus test was significant, but not of the individual tests were. Alternatively, among the sites that were identified by at least one of the pairwise tests, 85.2% of them are also identified by the omnibus test; for the remainder of the sites, the omnibus test is not significant. Among those sites, 89.6% had a single pairwise significant test (median omnibus corrected *p* = 0.103, IQR, [0.072*−*0.159]), 9.5%had two pairwise significant tests (0.066[0.056−0.081]), and 0.9% – three pairwise significant tests (0.058[0.053*−*0.071]).

To boost power in low information settings (small branch sets or low divergence), it may be advisable to run only the omnibus test, i.e. forego pairwise tests and the attendant FWER correction.

### Comparison with post-hoc tests

A reasonable heuristic approach for identifying sites that evolve differently between branch sets, *B*_1_ and *B*_2_ is to run an existing test which can determine whether the site evolved *non-neutrally* along either sets, and call the site differentially selected if there’s evidence of positive selection on one group but not another. Approaches like this have been commonly used in literature, e.g., Kapralov, Smith and Filatov (2012). One can also call a site differentially evolving if non-neutrality tests of *B*_1_ and *B*_2_ return *discordant results*. For example, *B*_1_ is negatively selected, but *B*_2_ is neutral, or *B*_2_ is positively selected and *B*_1_ is negatively selected. Contrast-FEL is, of course, a direct test of rate differences, so it could additionally identify, for instance, sites where *B*_1_ and *B*_2_ and both negatively/positively selected, but not at the same level. To illustrate the benefit of Contrast-FEL compared to the *post hoc approach* (which also requires at least two separate computational analyses, one for each branch set, so may be less computationally efficient), we performed post-hoc analyses based on independent FEL tests (one for each branch set) on a subset of 185070 sites from 425 alignments.

Using the LRT p-value cutoff of 5%, over all variable sites, Contrast-FEL achieves false positive rate (FPR) of 3.4%, power of 37.2%, positive predictive value (PPV) of 62.0% and negative predictive value of 91.1%. The “discordant results” post-hoc FEL approach, by comparison has FPR of 53.0%, power of 63.6%, PPV of 15.2% and NPV of 89.6%; the dramatic increase in the rate of false positives for the post hoc method is mostly (93.7%) due to cases, where a site that was simulated under the null is misclassified because one of the branch set if determined to be non-neutral by FEL and the other – neutral. All of the sites that were identified by Contrast-FEL, but not by the post hoc heuristic were those where FEL (correctly) determined that both branch sets were conserved, but the degrees of conservation, measured by the *β_i_/α<* 1 ratio were different. Empirical datasets analyzed in the following section provide concrete examples of such sites (they are in the CC FEL pattern).

The heuristic in which sites are called differentially evolving where only one the sets was under positive selection (the method commonly used in literature), has FPR of 3.3%, much reduced power of 17.1%, NPV of 94.6% and PPV of 25.9%.

### Empirical data

#### Drug resistance in HIV-1 reverse transcriptase (RT)

We applied Contrast-FEL to an alignment of 476 HIV-1 RT sequences with 335 codons isolated from 288 HIV-1 infected individuals, previously analyzed in (Murrell et al., 2012). There were two sequences sampled from each individual: one prior to treatment with reverse transcriptase inhibitors, and one following treatment. We partitioned the branches in the tree into three groups: pre-treatment terminal branches (*naive*), post-treatment terminal branches (*treated*), and the rest of the branches (nuisance group, Fig **??**.A). Because we expect the primary difference between the selective regimes on *naive* vs *treated* branches is due to the action of the antiretroviral drugs, most of the sites that have detectable differences in selective pressures should be implicated in conferring drug resistance. Using nominal p-value cutoff of 0.05, Contrast-FEL identifies 15 sites that evolve differentially, between which 12 are known drug resistance (DR) sites, achieving positive predictive value (PPV) of 0.8. Of the three non-DR sites that are found, codons 44 and 302 are actually more conserved in (lower *β*) in the treated group, which is a different mode of selective pressure than positive selection exerted by antiretroviral drugs. They are also not supported by the permutation test, which could indicate that these sites are picked up due to sampling variation. Among the 12 DR sites identified by LRT p-values, 10 are also supported by the permutation test – an indicator of robustness to branch sampling artifacts. The most conservative approach, based on FDR corrected q-values of 0.20 or lower, identifies 4 codons: 103, 181, 184 and 190, all of which are on the list of canonical escape mutations with very strong phenotypic effects (Rhee et al., 2003). All of these sites have many inferred mutations in the treated group, and large differences between inferred *β* rates, which places them in the large effect/large sample size category. As a comparison with one common practice to screen for differential selection today, we also used fixed effects likelihood (FEL, Kosakovsky Pond and Frost (2005)) to test each branch set for evidence of deviations from neutrality (either positive or negative selection). For site 190, subject to differential selection with a large effect size, FEL reveals that the treated branches experience positive selection (FEL *p ≤* 0.05), and naive branches – negative selection. However no other sites show such a clean pattern. For sites 103,151,184,215, test branches are subject to positive selection (FEL *p ≤* 0.05), while naive branches evolve neutrally. For sites 67,181,228, naive branches are subject to negative selection (FEL *p ≤* 0.05), while test branches evolve neutrally. For the remainder of the sites, neither branch class evolves in a way that is detectably different from neutrality according to FEL. This comparison highlights that comparing the results of two independent tests applied to subsets of the data to detect evolutionary differences is statistically suboptimal.

The performance of Contrast-FEL (a generalist method) in identifying sites of phenotypic relevance, compares favorably to the performance of a purpose-built MEDS method (Murrell et al., 2012), designed to find directional evolution along selected branches. MEDS identified 17 sites of which 10 were known DR sites (PPV of 58.9%, see Table 2 in Murrell et al. (2012), and both methods agreed on 9 sites. Of course, unlike MEDS, Contrast-FEL is not able to identify specific residues that may be the targets of selection at specific sites.

#### Selection on HIV-1 envelope conditioned on the route of transmission

We re-analyzed an alignment of 131 partial HIV-1 envelope (no variable loops) sequences with 806 codons from (Tully et al., 2016); these sequences were isolated from acute/early infections and represent “founder” viruses. These sequences were labeled by whether or not the infected individual was infected via a heterosexual (HSX) exposure, or men who have sex with men (MSM) exposure; interior branches were labeled as HSX or MSM if all of their descendants had the same label, and were left unlabeled (nuisance set) otherwise. Tully et al. (2016) found gene-wide differences in selection among branch groups (a larger proportion of sites, but subject to weaker positive selection, on HSX branches compared to MSM branches), using the RELAX test (Wertheim et al., 2015), but lacked the framework to pinpoint specific residues that evolved differentially. Contrast-FEL identified 32 differentially selected sites (p-value) of which 3 passed the FDR correction (Table 4). One of these sites, 626 is conserved in both branch sets, but at a stronger level (lower *β*) in the MSM set, while another (786), is positively selected in both, but at a stronger level on HSX branches.

**Table 4.** Sites evolving differentially between the HSX and the MSM sets in the HIV-1 env dataset from Tully et al. (2016). Other notation the same as in Table 3.

| Codon | $\alpha$ | $\beta$ (substitutions) | | p-value | q-value | Standard FEL p-value | | FEL pattern |
| --- | --- | --- | --- | --- | --- | --- | --- | --- |
|  |  | HSX | MSM |  |  | HSX | MSM |  |
| 49 | 0.56 | 0.12 (2) | 0.69 (12) | 0.0365* | 1.000 | 0.151(-) | 0.742(+) | NN |
| 50 | 0.30 | 0.61 (6) | 0.00 (2) | 0.0009* | 0.177 | 0.331(+) | 0.015(-) | NC |
| 53 | 0.88 | 0.00 (2) | 0.32 (9) | 0.0391 | 1.000 | 0.002(-) | 0.088(-) | CN |
| 142 | 4.13 | 5.09 (34) | 2.86 (37) | 0.0497 | 1.000 | 0.373(+) | 0.338(-) | NN |
| 158 | 0.62 | 2.24 (9) | 0.84 (10) | 0.0168 | 0.905 | 0.010(+) | 0.582(+) | PN |
| 197 | 1.88 | 2.93 (20) | 1.17 (23) | 0.0053* | 0.616 | 0.361(+) | 0.287(-) | NN |
| 233 | 0.20 | 0.25 (3) | 0.00 (1) | 0.0337* | 1.000 | 0.836(+) | 0.046(-) | NC |
| 264 | 0.33 | 1.91 (11) | 0.65 (9) | 0.0107 | 0.859 | 0.005(+) | 0.427(+) | PN |
| 275 | 2.45 | 4.16 (28) | 1.71 (28) | 0.0031 | 0.498 | 0.128(+) | 0.344(-) | NN |
| 303 | 1.63 | 6.98 (34) | 3.55 (39) | 0.0093* | 0.837 | 0.000(+) | 0.082(+) | PN |
| 336 | 3.80 | 1.09 (11) | 2.47 (35) | 0.0254 | 1.000 | 0.009(-) | 0.297(-) | CN |
| 344 | 0.40 | 1.25 (8) | 0.31 (7) | 0.0074* | 0.742 | 0.042(+) | 0.682(-) | PN |
| 408 | 0.52 | 0.70 (4) | 1.80 (22) | 0.0436 | 1.000 | 0.663(+) | 0.007(+) | NP |
| 442 | 0.00 | 0.31 (1) | 0.00 (0) | 0.0362* | 1.000 | 0.031(+) | 1.000(-) | PN |
| 530 | 0.69 | 0.29 (2) | 0.00 (3) | 0.0359* | 1.000 | 0.305(-) | 0.002(-) | NC |
| 572 | 1.55 | 4.01 (29) | 2.27 (33) | 0.0469 | 1.000 | 0.046(+) | 0.456(+) | PN |
| 574 | 1.10 | 0.52 (6) | 0.10 (5) | 0.0389 | 1.000 | 0.230(-) | 0.002(-) | NC |
| 598 | 0.59 | 1.13 (9) | 0.28 (7) | 0.0410 | 1.000 | 0.235(+) | 0.255(-) | NN |
| 626 | 12.77 | 4.14 (37) | 1.40 (41) | 0.0003* | 0.120 | 0.001(-) | 0.000(-) | CC |
| 672 | 2.46 | 3.15 (25) | 1.46 (37) | 0.0139* | 0.799 | 0.433(+) | 0.079(-) | NN |
| 683 | 1.88 | 2.05 (15) | 0.85 (17) | 0.0404* | 1.000 | 0.786(+) | 0.039(-) | NC |
| 685 | 0.00 | 1.16 (9) | 0.12 (2) | 0.0008* | 0.213 | 0.001(+) | 0.259(+) | PN |
| 690 | 1.24 | 0.28 (5) | 1.33 (18) | 0.0176* | 0.837 | 0.050(-) | 0.983(+) | CN |
| 702 | 0.00 | 1.13 (8) | 0.28 (4) | 0.0174* | 0.877 | 0.002(+) | 0.101(+) | PN |
| 703 | 1.31 | 1.27 (7) | 0.13 (11) | 0.0119* | 0.739 | 0.958(-) | 0.003(-) | NC |
| 720 | 0.49 | 1.47 (11) | 0.44 (13) | 0.0107 | 0.785 | 0.026(+) | 0.860(-) | PN |
| 722 | 0.00 | 0.13 (1) | 0.78 (9) | 0.0325 | 1.000 | 0.288(+) | 0.007(+) | NP |
| 734 | 0.50 | 2.23 (14) | 0.75 (10) | 0.0111 | 0.749 | 0.005(+) | 0.536(+) | PN |
| 773 | 0.25 | 0.00 (0) | 0.31 (7) | 0.0406 | 1.000 | 0.093(-) | 0.778(+) | NN |
| 781 | 0.51 | 0.50 (5) | 0.00 (2) | 0.0031* | 0.416 | 0.975(-) | 0.005(-) | NC |
| 786 | 1.40 | 6.67 (24) | 3.33 (38) | 0.0274 | 1.000 | 0.000(+) | 0.009(+) | PP |
| 804 | 0.25 | 0.00 (0) | 1.53 (15) | 0.0000* | 0.031 | 0.119(-) | 0.002(+) | NP |

#### Cell shape in epidermal leaf trichomes

Mazie and Baum (2016) investigated which codons in a developmentally important gene (BRT) in Brassicaceae (58 sequences, 318 codons) may be associated with the evolution of a different trichome cell shape in the genus *Physaria*. Using gene-level mean differences in *dN/dS* between subsets of branches, they identified that the average strength of selection is different in *Physaria* compared to the rest of the taxa. They then used a restricted branch-site (Yang and Nielsen, 2002) method to detect 10 codons that were subject to positive selection in the *Physaria* clade and 4 codons were “distinctive” (majority amino-acid was different in *Physaria*), but not positively selected. Contrast-FEL identified 29 differentially selected codons at *p ≤* 0.05 (18 at *q ≤.*20), including all 10 positive codons from Mazie and Baum (2016) and one out of four “distinctive” codons (Table 5). Given the general conservative nature of Contrast-FEL, it is reasonable to assume that it is more powerful (rather than prone to making more Type I errors) than the original test which was limited to a 50% subset of branches and used much more stringent parametric assumptions on rate distributions.

**Table 5.** Sites evolving differentially between *Physaria* and other taxa in the BLT gene from Mazie and Baum (2016). ^†^ – codon identified as positively selected in *Physaria* (Table 3 of Mazie and Baum (2016)), ^‡^ – codon identified as “distinctive” in *Physaria* (Table 4 of Mazie and Baum (2016)). Other notation the same as in Table 3.

| Codon | $\alpha$ | $\beta$ (substitutions) | | p-value | q-value | Standard FEL p-value | | FEL pattern |
| --- | --- | --- | --- | --- | --- | --- | --- | --- |
|  |  | *Physaria* | Other |  |  | *Physaria* | Other |  |
| 4 | 0.00 | 80.00 (2) | 0.76 | 0.0320 | 0.392 | 0.018(+) | 0.220(+) | PN |
| 43 | 0.69 | 4.27 (1) | 0.00 | 0.0110* | 0.194 | 0.251(+) | 0.043(-) | NC |
| 60 | 0.00 | 57.88 (1) | 0.82 | 0.0265 | 0.421 | 0.020(+) | 0.140(+) | PN |
| 107 | 0.93 | 1.70 (3) | 0.16 | 0.0123* | 0.206 | 0.511(+) | 0.108(-) | NN |
| 122 | 0.21 | 0.00 (0) | 1.40 | 0.0308* | 0.426 | 0.476(-) | 0.094(+) | NN |
| 126 | 1.62 | 0.89 (2) | 0.08 | 0.0467 | 0.550 | 0.563(-) | 0.017(-) | NC |
| 153<sup>†</sup> | 0.24 | 4.17 (6) | 0.23 | 0.0001* | 0.006 | 0.001(+) | 0.960(-) | PN |
| 156 | 0.66 | 2.91 (7) | 0.67 | 0.0283* | 0.409 | 0.051(+) | 0.986(+) | NN |
| 163 | 0.35 | 1.31 (2) | 0.12 | 0.0473 | 0.538 | 0.276(+) | 0.476(-) | NN |
| 164 | 0.32 | 2.04 (3) | 0.25 | 0.0267 | 0.404 | 0.088(+) | 0.847(-) | NN |
| 167<sup>†</sup> | 0.37 | 6.54 (10) | 0.70 | 0.0000* | 0.003 | 0.001(+) | 0.549(+) | PN |
| 169 | 5.04 | 0.74 (4) | 0.07 | 0.0476 | 0.522 | 0.016(-) | 0.000(-) | CC |
| 171<sup>†</sup> | 2.03 | 4.00 (4) | 0.00 | 0.0000* | 0.003 | 0.320(+) | 0.000(-) | NC |
| 173<sup>†</sup> | 0.47 | 4.12 (6) | 0.36 | 0.0011* | 0.044 | 0.026(+) | 0.820(-) | PN |
| 174<sup>†</sup> | 0.00 | 11.60 (8) | 0.31 | 0.0000* | 0.000 | 0.000(+) | 0.346(+) | PN |
| 175<sup>†</sup> | 0.85 | 3.67 (7) | 0.00 | 0.0000* | 0.002 | 0.046(+) | 0.006(-) | PC |
| 176 | 0.56 | 1.22 (1) | 0.00 | 0.0057* | 0.130 | 0.450(+) | 0.025(-) | NC |
| 178 | 1.29 | 1.93 (3) | 0.12 | 0.0085 | 0.160 | 0.636(+) | 0.026(-) | NC |
| 179 | 0.77 | 1.85 (2) | 0.00 | 0.0008* | 0.038 | 0.306(+) | 0.008(-) | NC |
| 180<sup>†</sup> | 0.17 | 2.22 (3) | 0.00 | 0.0001* | 0.006 | 0.008(+) | 0.161(-) | PN |
| 187 | 0.46 | 0.91 (2) | 0.00 | 0.0066* | 0.141 | 0.503(+) | 0.023(-) | NC |
| 188 | 1.31 | 1.03 (3) | 0.00 | 0.0075* | 0.149 | 0.790(-) | 0.001(-) | NC |
| 190<sup>†</sup> | 0.00 | 1.47 (4) | 0.07 | 0.0015* | 0.053 | 0.020(+) | 0.562(+) | PN |
| 191<sup>†</sup> | 0.45 | 2.49 (4) | 0.13 | 0.0025* | 0.080 | 0.046(+) | 0.302(-) | PN |
| 198 | 0.66 | 2.17 (2) | 0.00 | 0.0026* | 0.076 | 0.210(+) | 0.023(-) | NC |
| 255<sup>†</sup> | 2.06 | 1.47 (3) | 0.07 | 0.0047* | 0.124 | 0.636(-) | 0.000(-) | NC |
| 262 | 1.43 | 1.27 (3) | 0.17 | 0.0320* | 0.407 | 0.872(-) | 0.009(-) | NC |
| 270 | 2.32 | 0.87 (3) | 0.00 | 0.0047* | 0.115 | 0.248(-) | 0.000(-) | NC |
| 278<sup>‡</sup> | 1.64 | 0.86 (1) | 0.00 | 0.0320* | 0.424 | 0.567(-) | 0.001(-) | NC |

#### Evolution of Rubisco in C3 vs C4 photosynthetic pathway plants

Several studies comparing evolutionary selective pressures on the *rbcL* gene in C3 and C4 plants have identified several sites that appear to be under positive selection in either C3 or C4 plants (Kapralov and Filatov, 2007; Kapralov, Smith and Filatov, 2012), as well as several others that have different targets for directional evolution based on the pathway (Parto and Lartillot, 2018). In this alignment of 179 sequences and 447 codons, Contrast-FEL identified 15 sites that evolve differentially between C3 and C4 plants (LRT *p ≤* 0.05), of which 6 had been previously identified as being subject to differential directional selection by a mutation-selection model, and 5 additional sites were identified by this model (cf Table 6). An interesting example in this dataset is site 309 which was found as positively selected previously, but is classified as conserved in both C3 and C4 plants by FEL; this appears to be a result of the high synonymous rate inferred at the site, which is a hallmark false positive for standard selection analyses that ignore site-to-site synonymous rate variation (Kosakovsky Pond and Muse, 2005; Wisotsky et al., 2020). However, a weaker extent of conservation in C4 plants is inferred by Contrast-FEL at this site.

**Table 6.**
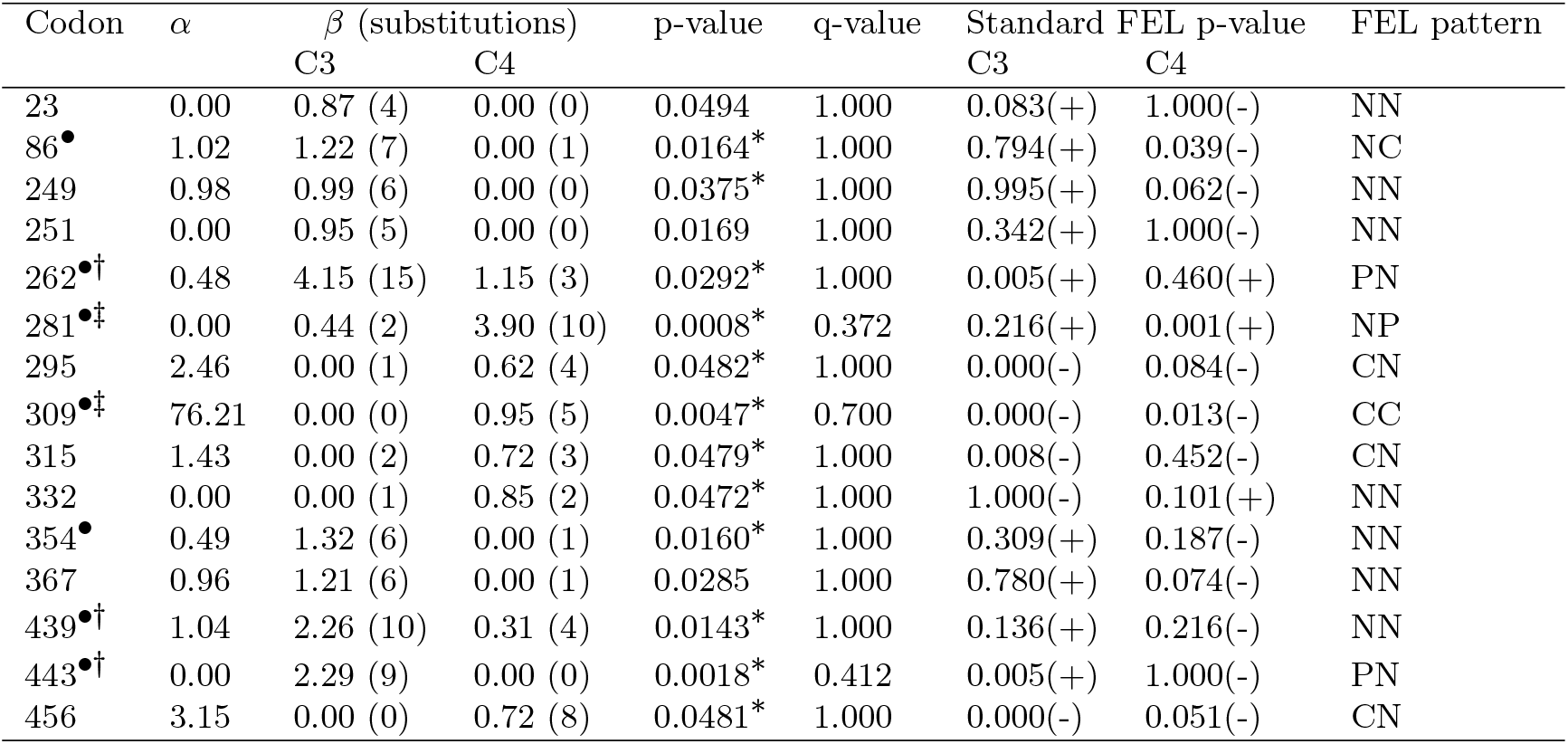
Sites evolving differentially between C3 and other taxa in the *r* bcL gene. ^†^ – codon identified as positively selected in C3 plants (^‡^ – C4) plants previously (Table 1 of Parto and Lartillot (2018)). • codon reported as differentially evolving by mutation-selection directional DS3 model in Parto and Lartillot (2018). Other notation the same as in Table 3.

| Codon | $\alpha$ | $\beta$ (substitutions) | | p-value | q-value | Standard FEL p-value | | FEL pattern |
| --- | --- | --- | --- | --- | --- | --- | --- | --- |
|  |  | C3 | C4 |  |  | C3 | C4 |  |
| 23 | 0.00 | 0.87 (4) | 0.00 (0) | 0.0494 | 1.000 | 0.083(+) | 1.000(-) | NN |
| 86<sup>•</sup> | 1.02 | 1.22 (7) | 0.00 (1) | 0.0164* | 1.000 | 0.794(+) | 0.039(-) | NC |
| 249 | 0.98 | 0.99 (6) | 0.00 (0) | 0.0375* | 1.000 | 0.995(+) | 0.062(-) | NN |
| 251 | 0.00 | 0.95 (5) | 0.00 (0) | 0.0169 | 1.000 | 0.342(+) | 1.000(-) | NN |
| 262<sup>•†</sup> | 0.48 | 4.15 (15) | 1.15 (3) | 0.0292* | 1.000 | 0.005(+) | 0.460(+) | PN |
| 281<sup>•‡</sup> | 0.00 | 0.44 (2) | 3.90 (10) | 0.0008* | 0.372 | 0.216(+) | 0.001(+) | NP |
| 295 | 2.46 | 0.00 (1) | 0.62 (4) | 0.0482* | 1.000 | 0.000(-) | 0.084(-) | CN |
| 309<sup>•‡</sup> | 76.21 | 0.00 (0) | 0.95 (5) | 0.0047* | 0.700 | 0.000(-) | 0.013(-) | CC |
| 315 | 1.43 | 0.00 (2) | 0.72 (3) | 0.0479* | 1.000 | 0.008(-) | 0.452(-) | CN |
| 332 | 0.00 | 0.00 (1) | 0.85 (2) | 0.0472* | 1.000 | 1.000(-) | 0.101(+) | NN |
| 354<sup>•</sup> | 0.49 | 1.32 (6) | 0.00 (1) | 0.0160* | 1.000 | 0.309(+) | 0.187(-) | NN |
| 367 | 0.96 | 1.21 (6) | 0.00 (1) | 0.0285 | 1.000 | 0.780(+) | 0.074(-) | NN |
| 439<sup>•†</sup> | 1.04 | 2.26 (10) | 0.31 (4) | 0.0143* | 1.000 | 0.136(+) | 0.216(-) | NN |
| 443<sup>•†</sup> | 0.00 | 2.29 (9) | 0.00 (0) | 0.0018* | 0.412 | 0.005(+) | 1.000(-) | PN |
| 456 | 3.15 | 0.00 (0) | 0.72 (8) | 0.0481* | 1.000 | 0.000(-) | 0.051(-) | CN |

#### Selection on cytochrome B of Haemosporidians infecting different hosts

Pacheco et al. (2018) performed an in-depth evolutionary analysis of three mitochondrial genes from 102 Haemosporidian parasite species partitioned into four groups based on the hosts. The analysis concluded that the genes were subject to mostly purifying selection, with different gene-level strengths of selection established using RELAX. For example, in the cytochrome B gene (376 codons) which we re-analyze here, selection in the plasmodium infecting avian hosts clade was intensified relative to the plasmodium infecting primate/rodent hosts. Because this analysis contained more than two branch groups, Contrast-FEL conducted 7 tests per site – the omnibus test and six pairwise group comparison (Table 7). Overall, 28 sites showed evidence of differential selection with at least one test (FWER corrected), and 5 tests passed FDR correction for the omnibus test. For clarity, we did not consider FEL analyses on individual branch sets and only focused on Contrast-FEL inference. Twenty-two of 28 sites were detected by the omnibus and between one and three pairwise tests, while six sites were reported only by one of the pairwise tests, highlighting the additional resolution offered by these more focused tests. Patterns of differences at individual sites varied widely, with every possible pair being significantly different at least once. The simplest case (e.g., 160 and 179) is a significant discordance between two groups of branches. Another repeated pattern is when one group of branches stands apart from all others (e.g., 89 and 102).

**Table 7.** Sites evolving differentially among the 4 branch groups in the Cytochrome B mitochondrial gene of Haemosporidians from Pacheco et al. (2018), according to the omnibus test or at least one pairwise test at LRT corrected p-value of ≤ 0.05. Individual pair tests in the last column and codes as follows. HA: Haemoproteidae vs Avian Hosts (one site), HM: Haemoproteidae vs Mammalian Hosts (two sites), HL: Haemoproteidae vs Leucocytozoon (six sites) AM: Avian Hosts vs Mammalian Hosts (four sites), AL: Avian Hosts vs Leucocytozoon (eighteen sites), ML: Mammalian Hosts vs Leucocytozoon (twenty sites). Other notation the same as in Table 3.

| Codon | $\alpha$ | $\beta$ (substitutions) | | | | | p-value | q-value | Significant<br>Pairwise Tests |
| --- | --- | --- | --- | --- | --- | --- | --- | --- | --- |
|  |  | Haemoproteidae | Avian Hosts | Mammalian Hosts | Leucocytozoon | Other |  |  |  |
| 56 | 3.81 | 0.00 (2) | 0.00 (2) | 0.00 (3) | 0.75 (4) | 0.00 | 0.0465* | 0.795 | AL,ML |
| 89 | 0.94 | 0.00 (1) | 0.00 (3) | 0.00 (4) | 1.33 (5) | 0.00 | 0.0001* | 0.017 | HL,AL,ML |
| 102 | 1.66 | 0.00 (2) | 0.00 (2) | 0.23 (5) | 1.83 (6) | 0.42 | 0.0014* | 0.103 | HL,AL,ML |
| 150 | 0.79 | 0.00 (1) | 0.00 (2) | 1.27 (9) | 0.00 (2) | 0.56 | 0.0075* | 0.403 | AM,ML |
| 158 | 0.42 | 0.00 (0) | 0.00 (0) | 0.00 (2) | 1.30 (5) | 0.00 | 0.0207* | 0.518 | AL,ML |
| 160 | 1.25 | 0.94 (6) | 0.00 (1) | 0.08 (8) | 0.43 (2) | 0.75 | 0.0422 | 0.755 | HA |
| 179 | 0.89 | 0.48 (2) | 0.00 (0) | 0.25 (6) | 1.46 (6) | 1.10 | 0.0171* | 0.495 | AL |
| 182 | 1.43 | 0.00 (0) | 0.00 (0) | 0.00 (0) | 2.23 (9) | 0.42 | 0.0001* | 0.013 | HL,AL,ML |
| 183 | 1.80 | 0.00 (0) | 0.00 (3) | 0.08 (2) | 1.20 (4) | 0.54 | 0.0058* | 0.361 | HL,AL,ML |
| 186 | 0.00 | 0.22 (1) | 0.00 (0) | 0.17 (2) | 0.65 (3) | 0.16 | 0.3127* | 1.000 | AL |
| 193 | 1.53 | 0.00 (0) | 0.00 (5) | 0.00 (1) | 1.07 (5) | 0.00 | 0.0147* | 0.462 | AL,ML |
| 194 | 1.05 | 0.00 (0) | 0.00 (0) | 0.00 (1) | 0.49 (4) | 0.19 | 0.1347* | 1.000 | ML |
| 222 | 0.64 | 0.00 (0) | 0.00 (0) | 0.00 (0) | 0.70 (4) | 0.00 | 0.0246* | 0.514 | AL,ML |
| 223 | 1.45 | 0.48 (3) | 0.00 (0) | 0.00 (1) | 0.70 (6) | 0.00 | 0.0200* | 0.538 | AL,ML |
| 248 | 0.48 | 0.00 (0) | 0.10 (1) | 1.01 (9) | 0.22 (1) | 0.64 | 0.0573* | 0.937 | AM |
| 253 | 1.67 | 0.00 (4) | 0.00 (4) | 0.00 (6) | 0.95 (4) | 0.40 | 0.0132* | 0.452 | AL,ML |
| 256 | 1.92 | 0.00 (6) | 0.00 (3) | 0.73 (8) | 0.00 (0) | 0.64 | 0.0386* | 0.725 | AM |
| 283 | 0.38 | 0.00 (1) | 0.16 (1) | 0.00 (0) | 1.36 (6) | 0.64 | 0.0241* | 0.534 | ML |
| 285 | 1.17 | 0.22 (1) | 0.00 (1) | 0.39 (5) | 1.44 (6) | 1.01 | 0.0852* | 1.000 | AL |
| 289 | 3.28 | 0.00 (0) | 0.00 (1) | 0.00 (0) | 1.31 (3) | 0.00 | 0.0012* | 0.110 | HL,AL,ML |
| 309 | 1.05 | 0.83 (2) | 0.36 (6) | 1.11 (9) | 4.24 (8) | 0.61 | 0.0822* | 1.000 | AL |
| 310 | 1.96 | 0.22 (1) | 0.00 (1) | 0.00 (2) | 0.83 (4) | 0.00 | 0.0119* | 0.499 | AL,ML |
| 331 | 0.15 | 0.00 (0) | 0.00 (0) | 0.00 (0) | 7.31 (9) | 0.00 | 0.0000* | 0.012 | HL,AL,ML |
| 338 | 0.00 | 0.00 (0) | 0.00 (0) | 0.00 (0) | 0.98 (3) | 0.00 | 0.0115 | 0.541 | AL,ML |
| 341 | 0.43 | 0.49 (3) | 0.28 (2) | 0.00 (2) | 0.83 (6) | 0.45 | 0.1118* | 1.000 | ML |
| 343 | 0.25 | 0.97 (3) | 0.14 (1) | 0.00 (1) | 1.23 (4) | 1.05 | 0.0224* | 0.527 | HM,ML |
| 351 | 1.75 | 0.00 (2) | 0.00 (3) | 0.91 (8) | 0.88 (4) | 0.25 | 0.0367* | 0.727 | AM |
| 366 | 0.52 | 1.43 (5) | 0.20 (3) | 0.00 (1) | 0.61 (6) | 0.67 | 0.0123* | 0.463 | HM,ML |

## Discussion

To narrow down the genetic basis of adaptation, many studies contrast evolution between different subsets of branches in a phylogenetic tree, selected to represent different phenotypes, environments, or fitness regimes. As we sequence more organisms and obtain better annotations of function and phenotypic differences, such contrast analyses are likely to become more common. However, with a few exceptions, methods that researchers have adopted to find differentially evolving sites were not designed to explicitly test for such differences. The Contrast-FEL approach, presented here, addressed this methodological shortcoming, and establishes a formal statistical framework for testing for differences in evolutionary rates among two or more sets of branches at individual sites. Unlike approaches which infer something about each branch set separately (e.g., is branch set X under positive selection at site S?), and compare these inferences in a *post hoc* fashion (site S is selected on branch set X but not on set Y), Contrast-FEL enables direct rigorous testing for differences in non-synonymous evolutionary rates at individual alignment sites in predefined non-overlapping sets of branches and is computationally feasible for all but the largest comparative analyses. Contrast-FEL has good operating characteristics on data simulated under a broad range of scenarios, and finds large numbers of differentially evolving sites in empirical datasets. When prior results are available, we find that in addition to recovering many sites with known effect on phenotype (e.g., HIV-1 drug resistance), or sites found with alternative methods (*rbcL*, BRT), Contrast-FEL reports subtler differences, such as sites that are subject to differing degrees of conservation, or not subject to detectable positive or negative selection in either subset. Therefore, Contrast-FEL may enable more precise and powerful comparative analyses, and it is also the first method of this class that is able to compare selective pressures among more than two groups of branches. The general framework of site-level rate comparison using likelihood ratio tests presented here can be readily extended to compare other types of evolutionary parameters, e.g., rates that are informed by amino-acid properties (Delport et al., 2010), rates of instantaneous substitutions that involve multiple nucleotides (Venkat, Hahn and Thornton, 2018), or rates that influence positional synonymous substitutions (Rubinstein et al., 2011)

As with any statistical inference method, it is important to appreciate when it will work well and when it will not. Since Contrast-FEL tests for significant differences in *dN/dS* rates, a positive result means that the sites are subject to different selection intensities, e.g. purifying vs neutral or positive diversifying, and not that they evolve towards different target residues (directional selection). It will work best on sufficiently large alignments with multiple substitutions in each of the branch sets, while, on small or genetically similar datasets, power will be low for all but the most dramatic differences. Our simulations provide guidelines for performance, and, if desired, power simulations for a specific dataset can be used to determine what effect size can be realistically detected. Since it is reasonable to assume that in most alignments most sites do not evolve differentially (or at least not dramatically so) along subsets of branches, care must be taken to control for false discovery. Our empirical data analyses on alignments with prior site-level results do not indicate a dramatically inflated rate of false positives (the lists of sites overlap substantially with those identified previously). On the other hand, being too conservative will lead to loss of power, since for site-level inference it does not grow with the length of the alignment (Scheffler, Murrell and Kosakovsky Pond, 2014). Depending on the desired tradeoffs between precision and recall, the user may choose to use site-level p-values (highest power, lowest precision), permutation p-values (intermediate power and precision), or q-values (lowest recall, highest precision), each of which is available in the HyPhy (http://www.hyphy.org) implementation of Contrast-FEL.

**FIG 1.**
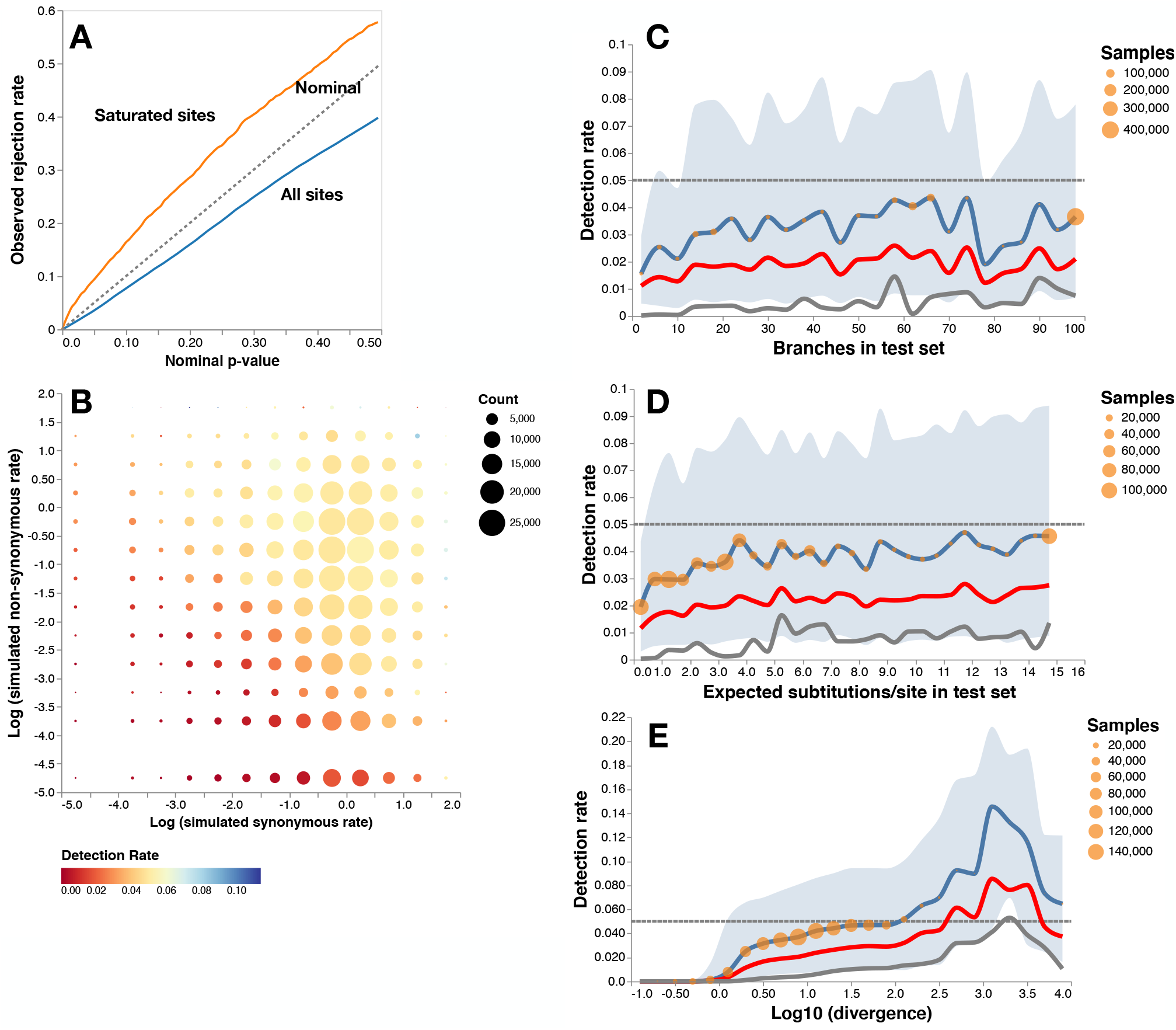
Contrast-FEL performance on null data (error control). The plots are based on 1090929 variable sites simulated with equal non-synoymous rates on two branch sets (see text for simulation details). **A.** Q-Q plots of LRT p-values for all sites (blue line) and for 3684 *saturated* sites (log10 of the divergence level between 2.5 and 3.5, orange line). **B.** Detection rate as a function simulated synonymous and non-synonymous substitution rates (log10 scale). **C.** Detection rate as a function of the number of branches in the test set (binned in increments of 5). Blue line: proportion of sites with LRT *p<* 0.05, red line: proportion of sites with permutation *p<* 0.05, grey line: proportion of sites with *q <* 0.20. Blue area plot shows for the proportion of sites with LRT *p<* 0.01 (lower) and LRT *p<* 0.1 (upper). Orange circles reflect the number of sites contributing to each bin. **D.** Detection rate as a function of the total branch length of the test set of branches (binned in increments of 0.5); same notation as in (C) otherwise. **E.** Detection rate as a function of the log10 of the divergence level at the site (binned in increments of 0.25); same notation as in (C) otherwise.

**FIG 2.**
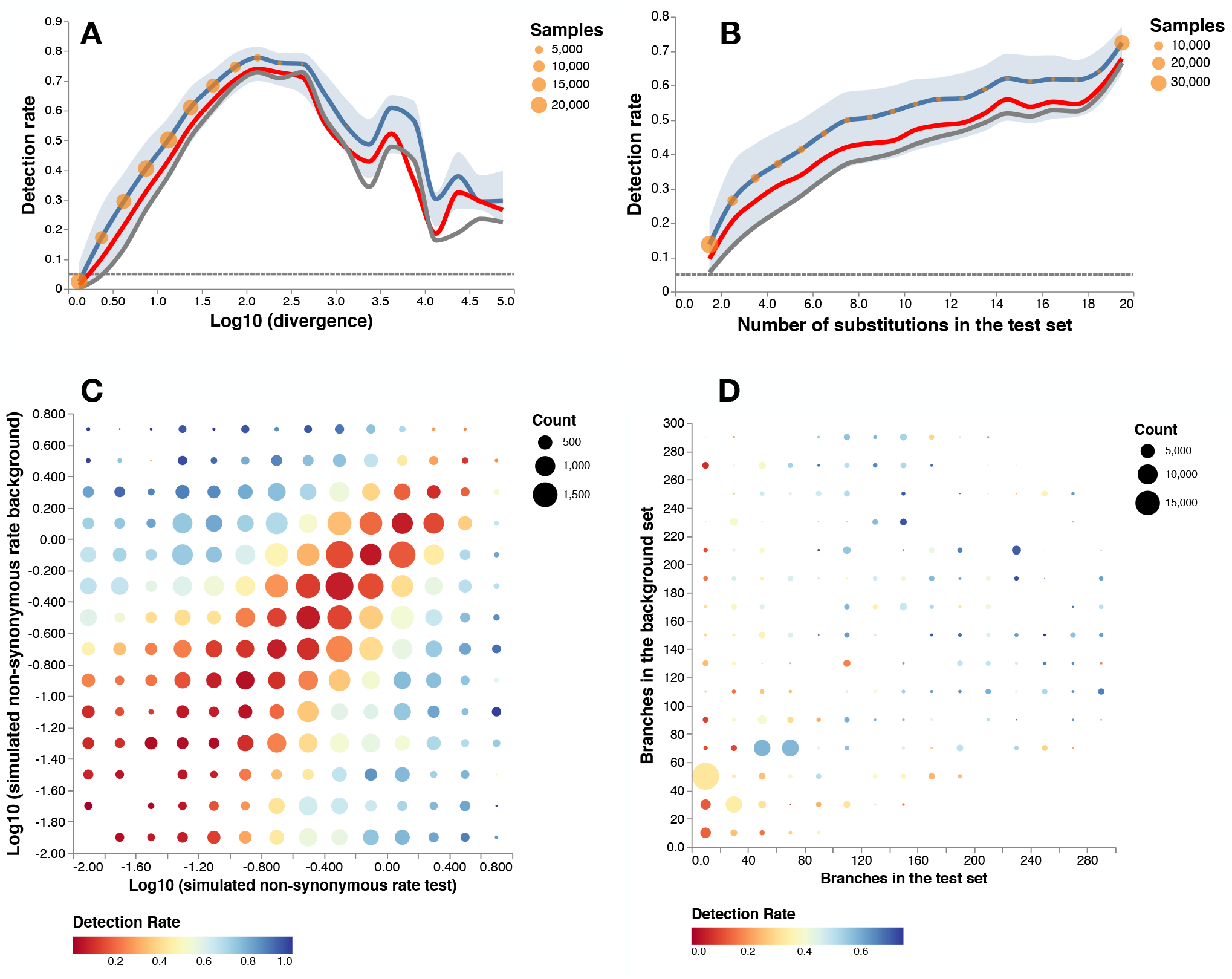
Contrast-FEL performance data with rate differences (power). The plots are based on 139753 variable sites simulated with unequal non-synoymous rates on two branch sets (see text for simulation details). **A.** Detection rate as a function of the log10 of the divergence level at the site. Blue line: proportion of sites with LRT *p<* 0.05, red line: proportion of sites with permutation *p<* 0.05, grey line: proportion of sites with *q <* 0.20. Blue area plot shows for the proportion of sites with LRT *p<* 0.01 (lower) and LRT *p<* 0.1 (upper). Orange circles reflect the number of sites contributing to each bin. **B**. Detection rate as a function of the number of inferred substitutions in the test set; same notation as in (A) otherwise. **C** Detection rate as a function of the the simulated non-synonymous rates in test and background branch sets and, **D.** the numbers of branches in the test and reference set.

**FIG 3.**
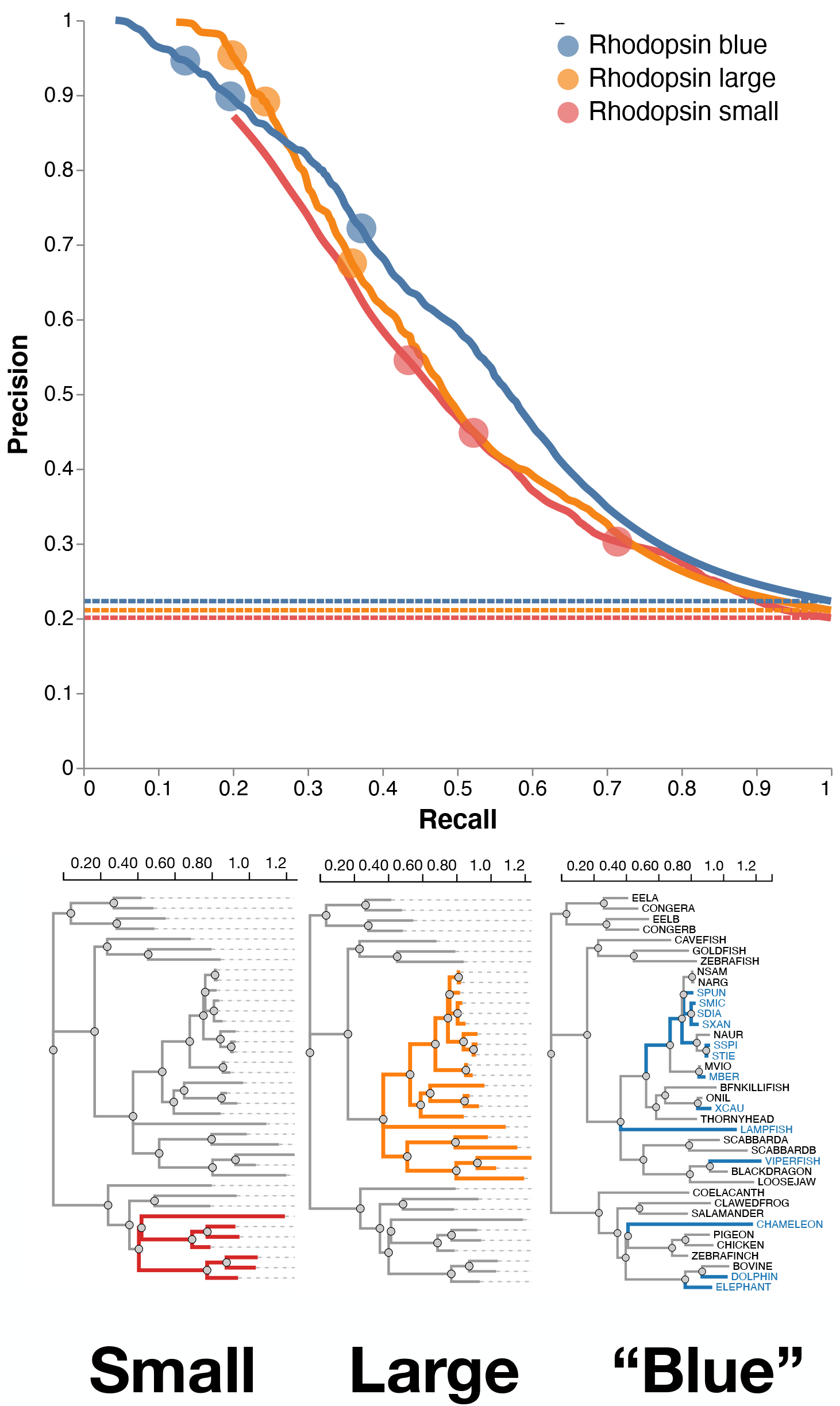
Contrast-FEL performance on vertebrate rhodopsin simulations. Precision-recall curves for the three sets of simulations, all based on the vertebrate rhodopsin tree from Yokoyama et al. (2008), with different choices for the “test” branch set. Dotted lines show corresponding base rates for “no-skill” classifiers in each case (i.e., classify all sites as differentially selected). Circles on the individual curves show (left-to-right) precision recall values for *q* = 0.1*,q* = 0.2*,q* = 0.5. There were a total of 37565 variable sites for the “small” case, 15010 sites for the “large” case, and 37401 sites for the “blue” case.

**FIG 4.**
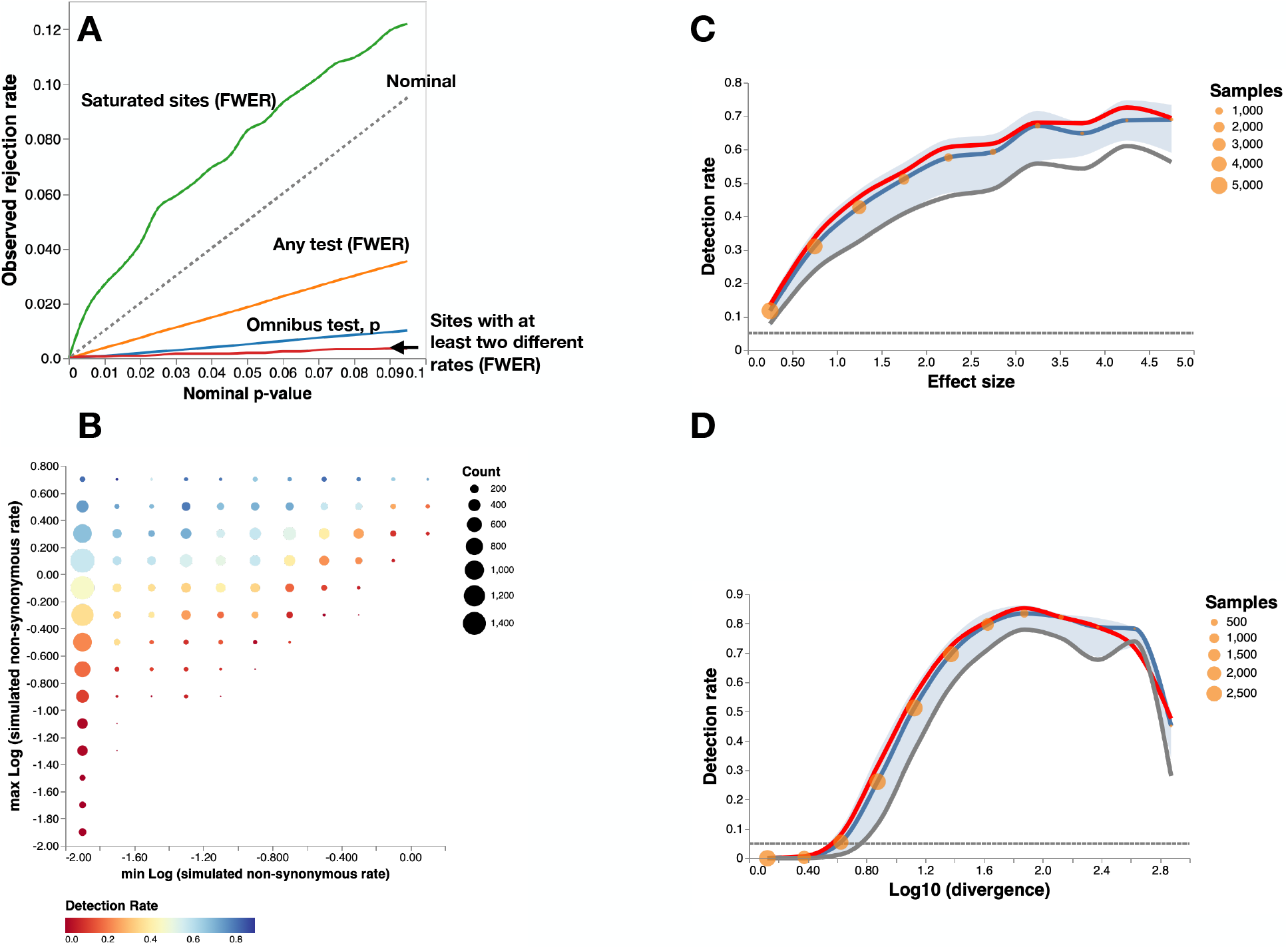
Contrast-FEL performance data with rate differences using four branch sets. **A.** Q-Q plots of either omnibus test p-values (blue) or family-wise error rates (FWER, orange), which are based on rejections of any of the true nulls among 151838 sites simulated where all branches had the same non-synonymous rate. Green line shows the Q-Q plot of the FWER rates on 1944 *saturated* sites (log10 of the divergence level above 2.5). Red line shows the FWER rate for the 3702 datasets where some, but not of the nulls were true (i.e., some branch sets shared rates, while others did not). **B.** Detection rate as a function of the log10 of the lowest and highest non-synonymous rates (rates lower than 0.01 are shown as 0.01) computed on 18141 sites where at least two rates were different. **C.** Detection rate as a function of the effect size, measured as the maximum difference between non-synonymous rates among branch classes. Blue line: proportion of sites with LRT *p<* 0.05 (the omnibus test), red line: proportion of sites with permutation *p<* 0.05, grey line: proportion of sites with *q <* 0.20. Blue area plot shows for the proportion of sites with LRT *p<* 0.01 (lower) and LRT *p<* 0.1 (upper). Orange circles reflect the number of sites contributing to each bin. **D.** Detection rate as a function of the log10 of the divergence level at the site

## Acknowledgments

This work was supported in part by grants R01 GM093939 (NIH/NIGMS) and AI134384 (NIH/NIAID).

## Supplementary Material

**FIG. S1.**
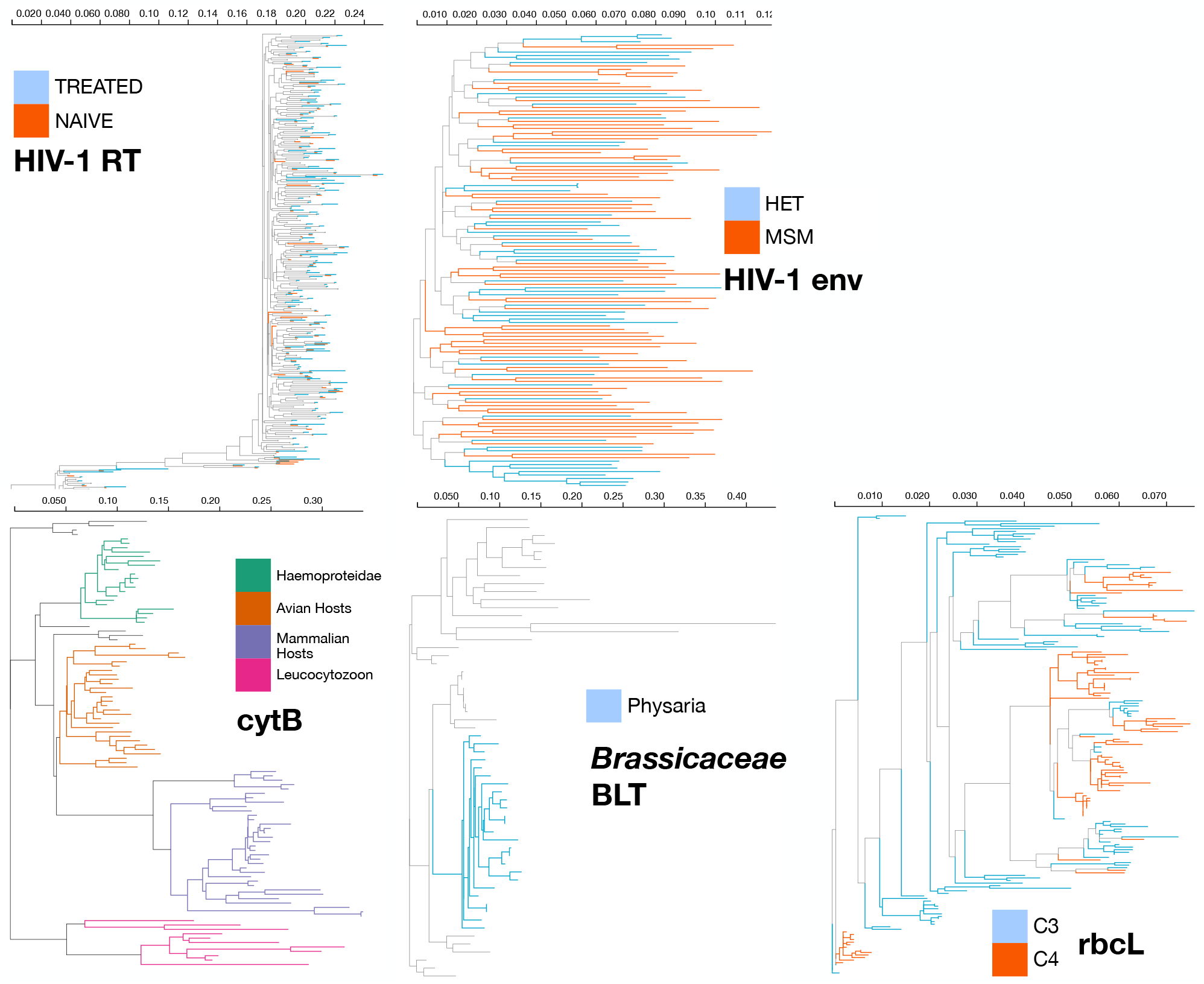
Trees and branch partitions for the empirical datasets. Branches shown in gray are in the nuisance (background) class. Alignments and trees can be downloaded from data.hyphy.org/web/contrast-FEL/.

